# MPFIL: Multi-Level Protein Feature Integration for Predicting Mutation-Induced Protein-Protein Binding Affinity Changes

**DOI:** 10.64898/2026.10.22.760922

**Authors:** Hao Zhang, Yang Liu, Limin Yu, Maozu Guo

## Abstract

—Predicting mutation-induced changes in protein-protein binding affinity (ΔΔ*G*) is challenging because mutations can affect both local residue environments and the structural organization of protein complexes. Existing approaches often rely on handcrafted physicochemical, energetic, or evolutionary features, which introduce additional computational costs and require domain-specific knowledge. In this work, we propose MPFIL, a multi-level protein feature integration framework for predicting mutation-induced protein-protein binding affinity changes. MPFIL integrates structural and sequential information to characterize mutation effects from complementary perspectives. Its structural feature extraction module captures mutation-induced structural changes and differential information between wild-type and mutant complexes, while sequence feature extraction modules model local residue context and long-range dependencies. The resulting features are integrated for ΔΔ*G* prediction. We evaluate MPFIL on three benchmark datasets. Experimental results demonstrate competitive and consistent performance across different protein complex settings, while ablation studies verify the contribution of the structural and sequential features. These results support the effectiveness of multi-level protein feature integration for modeling mutation-induced changes in protein-protein binding affinity.

## I. Introduction

Protein-protein interactions (PPIs) are fundamental to many biological processes, including cellular regulation, enzyme catalysis, and gene expression [1]. Mutations can alter protein structures and intermolecular interactions, thereby affecting the binding affinity, specificity, and stability of protein complexes [2]. Accurate prediction of mutation-induced binding affinity changes (ΔΔ*G*) is therefore important for understanding mutation effects and supporting applications such as disease mechanism analysis and therapeutic development.

Experimental characterization of mutation effects is often time-consuming and costly, motivating the development of computational approaches. Early machine learning (ML)-based methods typically incorporate physicochemical, energetic, and evolutionary features to characterize mutation effects. For example, mmCSM-PPI [3] uses features from multiple categories, including kinetics, residue protection, and residue environmental properties. With the increasing availability of protein structures, deep learning (DL)-based approaches have further explored automated feature extraction. Top-NetTree [4] combines convolutional neural networks (CNNs) with gradient boosting trees (GBT), while PerSpectEL [5] employs ensemble learning for protein representation. More recently, graph neural networks (GNNs) have been introduced to model structural information in protein complexes. GeoPPI [6] uses a geometric encoder to capture atomic-level structural features, and SFM-Net [7] integrates GNN-based and evolutionary features for prediction.

In order to describe the mutation effects, most of these methods involve a substantial number of complex biological features, such as physicochemical properties, energy parameters, and evolutionary information, which typically lead to an increase in computational and processing cost. The effectiveness of the methods is likewise highly dependent on expert knowledge, and the development will be limited in the face of the rapid growth of available protein data and the deficiency of expert knowledge data. Furthermore, despite the existence of prior studies that have employed encoders for protein structures, these methods lack consideration of information regarding structural changes induced by mutations, which is highly valuable for understanding the behavior and geometric regulations of protein complexes. The intrinsic relationships between mutation sites and neighboring residues in protein sequences still require further exploration. The design of solutions that tackle these limitations remains a significant challenge. We therefore consider constructing a more generalized protein feature extraction framework to address the aforementioned issues, which will obviate the introduction and processing of intricate biological features and expert prior knowledge.

In this paper, we propose MPFIL, a novel multi-level protein feature integration framework for predicting mutation-induced binding affinity changes in protein complexes, which enables the effective integration of informative structural features and multi-perspective sequential features. Specifically, we design a multi-perspective feature extraction network for protein sequences to fully exploit sequential information and the inherent contextual relationships between mutations and neighboring residues. Furthermore, the structural changes in protein complexes caused by mutations are crucial for understanding protein behavior and structural variations. To describe the structural changes and to better explore and utilize the structural information, we consider designing a GNN-based structural feature extraction network and the corresponding differential feature utilization method. The proposed framework integrates all-level extracted features with the objective of accurately predicting PPI ΔΔ*G*, while promoting a more comprehensive and systematic understanding of the properties and behaviors of protein complexes.

We conduct detailed evaluations of the proposed model on three benchmark datasets, including SKEMPI [8], SKEMPI-2.0 [9], and the membrane protein dataset MPAD [10]. The experimental results show that MPFIL achieves strong and consistent performance across these datasets, demonstrating its effectiveness and excellence for ΔΔ*G* prediction. Furthermore, we conduct ablation studies and distribution comparisons between the experimental and predicted values to verify the effectiveness of the proposed method in modeling and learning protein features. The contributions of our work can be summarized as follows:

1. We propose a novel multi-level protein feature integration framework for PPI ΔΔ*G* prediction that systematically integrates and explores diverse structure-sequence features, while avoiding reliance on complex biological features.
2. We design a graph-level encoding module for protein structure to accurately describe and capture mutation-induced structural changes and differential information between wild-type and mutant protein complexes.
3. We introduce various feature extraction modules to fully exploit multi-perspective sequential features, including the inherent contextual relationships between the mutation site and neighboring residues, global semantic relationships, and biological regulation.
4. We demonstrate the excellent performance of the proposed MPFIL method in comparison with several competitive methods for predicting ΔΔ*G*, via experiments on three benchmark datasets.

## II. Related Work

Research on mutation effect prediction has primarily focused on individual proteins, while predicting mutation-induced changes in the binding free energy of protein complexes remains more challenging due to their complex intermolecular interactions. Early approaches mainly relied on biophysical modeling and statistical potentials. For example, Li et al. [11] combine molecular mechanics force fields with statistical potentials for mutation effect prediction, while Delgado et al. [12] estimate free energy changes through energy optimization in biomolecular modeling. Although these approaches provide physically interpretable predictions, their computational cost and predictive performance can limit their practical application.

With the development of machine learning and deep learning, learning-based approaches have become increasingly prevalent. Wang et al. [4] combine convolutional neural networks (CNNs) and gradient boosting trees (GBT) with topological representations to predict affinity changes. Rodrigues et al. [3] employ multiple categories of biological features, including kinetics, residue protection, and residue environmental properties. Wee and Xia [5] combine CNN and GBT predictions using persistent spectral representations of PPIs. Xue et al. [13] incorporate deep residual recurrent CNNs with topological, structural, and functional information. More recently, Liu et al. [7] and Rimal et al. [14] have explored deep neural networks for general and membrane proteins, respectively, by incorporating molecular and biological features.

Despite these advances, existing methods still face several challenges. Many learning-based approaches depend on diverse handcrafted biological features, resulting in additional feature engineering and computational requirements. Sequence-based approaches may also have difficulty capturing sufficient structural information, while structure-based methods do not always explicitly model the structural differences induced by mutations. These limitations motivate MPFIL, which integrates structural and sequential features to characterize mutation effects without relying on extensive handcrafted biological descriptors.

## III. Framework

### A. Problem Formulation

Given the wild-type (*WT*) protein complex and the corresponding mutant-type (*ℳT*) protein complex, the mutation-induced binding affinity changes can be calculated as:

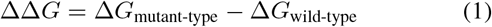

where Δ*G*_mutant-type_ and Δ*G*_wild-type_ represent the free energy difference between the folded and unfolded states of the wild-type complex and the mutant-type complex, respectively. The global objective function is defined as the application of a neural network *f* to approximate the solution to the mutation effects prediction problem as follows:

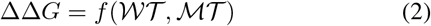

In this paper, the neural network *f* is defined as MPFIL, which is illustrated in Fig. 1, and details of the architecture are presented in the following subsections.

**Fig 1.**
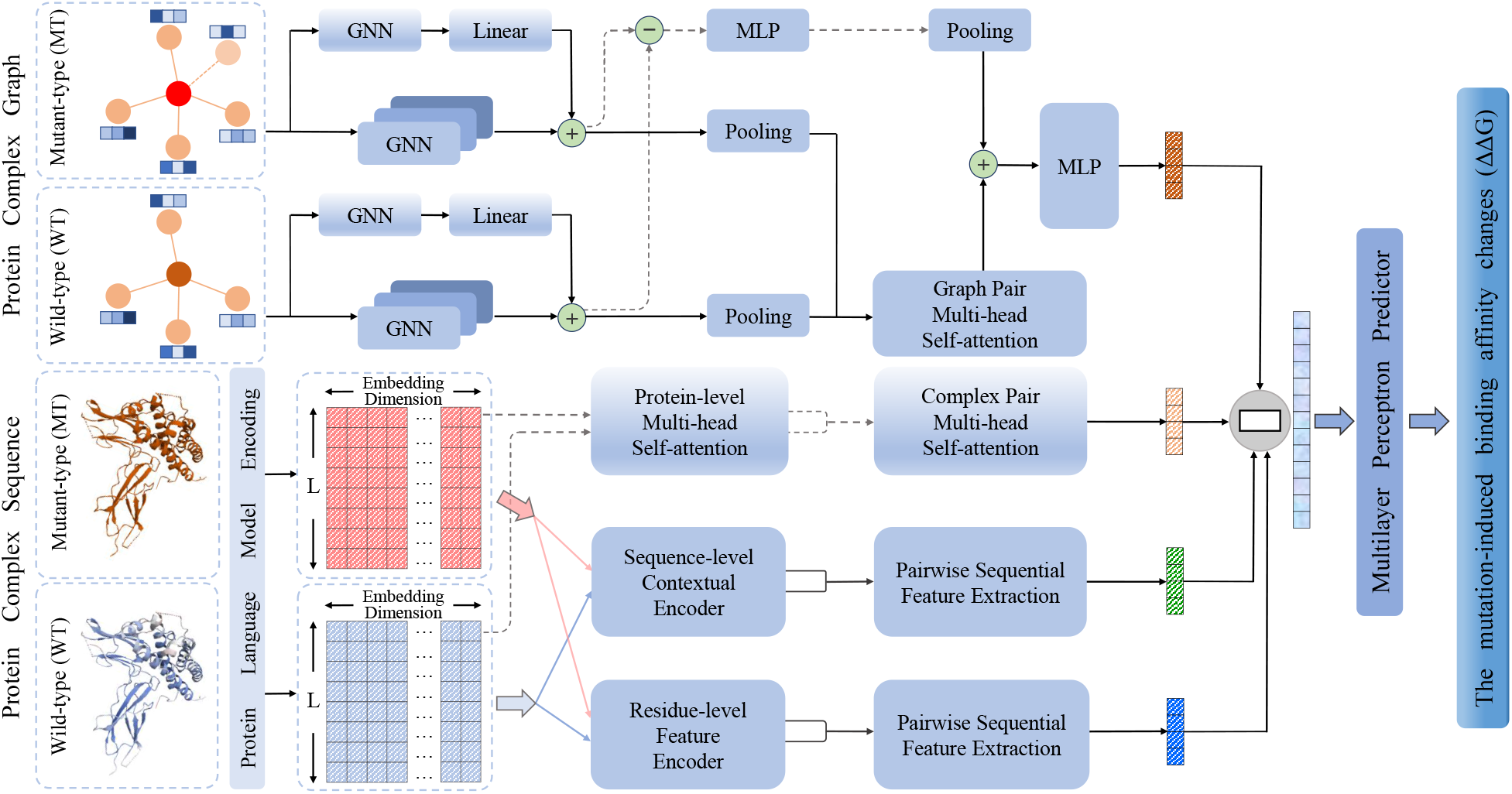
Overview of the proposed multi-level protein feature integrated learning framework(MPFIL)

### B. Protein Complex Representation

We represent protein complex mutations in two formats: the sequence and the undirected graph, facilitating comprehensive and efficient learning of protein features at multiple levels.

#### 1) Sequence Features

In the proposed method, the protein language model [15] functions as the input vector mapping layer, encoding the protein as sequential features. The objective of this process is to map proteins into high-dimensional embedding vector representations, which embody positional dependencies, implicit physicochemical signals, and syntactic features of the life language.

Given a protein chain *p* = *{a*_1_, *a*_2_, …, *a*_*L*_*}* from the protein complex, the protein language model encodes it into an informative embedding representation, in which each amino acid is transformed into a *d*-dimensional vector ***a***_*l*_, where 1 *≤ l ≤ L*, and *L* is the protein length. Each protein sequence is prefixed with a specific token, *⟨CLS⟩*, which is encoded to represent and capture the global level sequential feature to be further explored in subsequent modules. The embedding vector of the special token is denoted by ***a***_0_, and the vector representation of a protein is denoted by ***p*** = {***a***_0_, ***a***_1_, ***a***_2_, …, ***a***_*L*_*}*.

#### 2) Graph Features

In the graph representation, the mutation site is depicted as the central node, with the remaining nodes representing surrounding residues, and edges denoting interactions between the mutation site and other residues. As residues in closer proximity to the mutation site exert a more pronounced influence on local structure and conformation, residues within 10 A^*°*^ of the mutation site are considered when constructing edges for the central node [7]. The approach ensures that the information updates of GNN emphasize the central node, thereby highlighting the structural variation information most relevant to the mutation. The wild-type graph is constructed using spatial coordinates obtained from the Protein Data Bank (PDB) files. Mutation-induced structures are computed using FoldX [12], after which the resulting PDB files are used to construct the mutant-type graph.

The initial graph representation of a protein complex is defined by a node feature matrix and an adjacency matrix. The construction of node features is based on the embedding representation of the protein sequence. Based on the spatial coordinates, we calculate the Euclidean distances between the mutation site and other residues to construct a distance matrix. The values in the distance matrix are then compared with a predefined threshold to construct an adjacency matrix. Given a raw wild-type or mutant-type protein graph *G* with *n* nodes, we denote the node feature matrix as **V** *∈* ℝ^*n×d*^ and the adjacency matrix as **A** *∈* ℝ^*n×n*^.

### C. Graph-level Structural Feature Encoding

The graph-level structural feature encoding module aims to capture the mutation-induced structural changes and differential information in protein complexes, providing a more accurate description of the impact of mutations on the structure.

Given the wild-type graph *G*_*W*_ and the mutant-type graph *G* _*ℳ*_, both are fed into the GNN-based structural feature extraction module, which consists of three consecutive GNN layers with a residual GNN connection. The computation of the GNN-encoded feature is defined as follows:

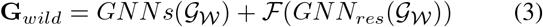

where **G**_*wild*_ ∈ ℝ^*n×d*^,*n* represents the number of nodes in the wild graph *G*_*W*_, ℱ denotes a linear layer, and *GNNs* and *GNN*_*res*_ denote the three stacked GNN layers and the residual GNN layer, respectively. The GNN-encoded feature representation **G**_*mut*_ of the mutant-type graph is obtained by following the same procedure. The process enhances the extraction of expressive and deep graph representations, enabling subsequent analysis of mutation-induced structural changes.

Subsequently, explicit and implicit operations are performed on the graph representations to capture structural differential information at various levels. Inspired by residual networks, the structural difference features of the complex graph pair are combined via element-wise summation. The resulting structural difference features are further aggregated and subsequently passed through a nonlinear layer and a multi-layer perceptron (MLP) to obtain the final encoded representation. The computational process is outlined as follows:

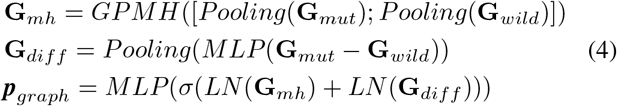

where *Pooling* denotes the global mean pooling operation, *GPMH* represents the multi-head self-attention block for the complex graph pair, *LN* denotes the layer normalization, *σ* denotes the activation function SiLU, and ***p***_*graph*_ ∈ ℝ^*d*^is the new graph-level structural representation of the complex pair.

The new representation learns and acquires more complex and enriched structural features of wild-type and mutant complexes, effectively integrating mutation-induced structural changes and differential information.

### D. Multi-perspective Sequential Feature Encoding

The multi-perspective sequential feature encoding component aims to fully exploit sequential information and inherent contextual relationships between mutations and neighboring residues.

#### 1) Global level Differential Feature

The global level differential feature encoding module aims to integrate full-sequence representations across molecular chains within a complex as well as between wild-type and mutant-type complexes, enabling comprehensive learning of the complete biological regulation of the complexes and thereby contributing to a deeper understanding of their differences.

Given protein chains *p*_1_ and *p*_2_ within the mutant-type complex, their vector representations of global sequential features ***a***_0_ and ã_0_ are concatenated and fed into the protein-level multihead self-attention block to compute a new representation of the complex, thereby modeling interactions between internal protein chains, as follows:

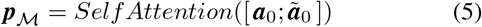

where ***a***_0_, ã_0_ ∈ ℝ^*d*^, and ***p***_*ℳ*_ ∈ ℝ^*d*^ is the representation of the mutant-type complex. The representation ***p***_*W*_ of the wild-type complex is obtained by following the same procedure.

The representations of the complexes are concatenated and processed through the complex pair multi-head self-attention block to obtain a new representation of the complex pair:

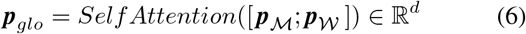

The new representation learns the complete biological regulation of wild-type and mutant-type complexes, and incorporates significant differential information between them.

#### 2) Full-sequence Context-level Feature

To capture broader contextual information beyond the binding interface, the sequence-level contextual encoder block encodes the complete amino acid sequences of protein chains within the complex. Given the full-length sequence features of a protein chain ***p*** ∈ ℝ^*L×d*^as input, the computation is performed as follows:

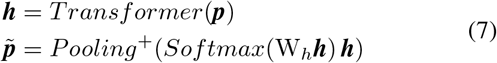

where ***h*** ∈ ℝ^*d*^ denotes the hidden representation output by the *Transformer* encoder, W_*h*_ ∈ ℝ^*d*^ is a trainable weight vector, *Pooling*^+^ denotes the sum pooling operation, and 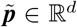 is the new representation of the protein chain that captures and incorporates the long-range interactions among protein chains.

In order to further compress features for subsequent prediction while integrating differential features between complex sequences, the context-level sequence feature representations 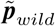 and 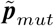 are jointly fed into the pairwise sequential feature extraction block. The block comprises convolutional layers and nonlinear layers, ultimately functioning to output the full-sequence context-level feature representation ***p***_*context*_ with 2 ×*d* dimensionality for the protein complex pair. Details of the extraction block architecture are provided in Figure 2. The module effectively captures and learns long-range dependencies, semantic relationships, and interactions among amino acid residues across the full-length protein complex sequence. The resulting context-level sequential feature representations complement features encoded at other levels.

**Fig 2.**
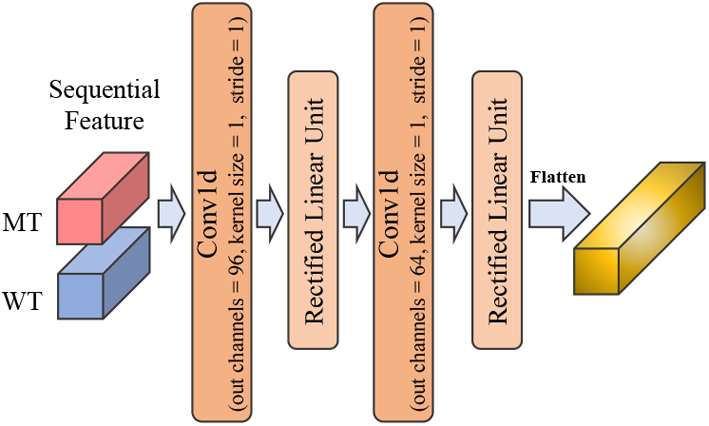
Pairwise sequential feature extraction block.

#### 3) Residue-level Sequential Feature

The residue-level feature encoder block employs multiple CNNs with varying kernel sizes, facilitating the detection and capture of local sequence feature patterns across multiple receptive fields, including short motifs, medium-range dependencies, and conserved fragments. Initially, a small-kernel convolution is applied to compress the channel dimension and model the interactions among proximal residues. Convolutions, non-linear activation layers, and max-pooling layers are then alternately employed to further extract crucial protein features and reduce the sequence length. The flattened output from the encoder produces a new vector with the same dimensionality as the input, i.e., 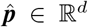. Details of the architecture are provided in Figure 3. The encoded features of the complex pair are integrated and fed into the pairwise sequential feature extraction block for further compression and refinement, resulting in a residue-level sequential feature representation ***p***_*residue*_ ∈ ℝ^2*×d*^.

**Fig 3.**
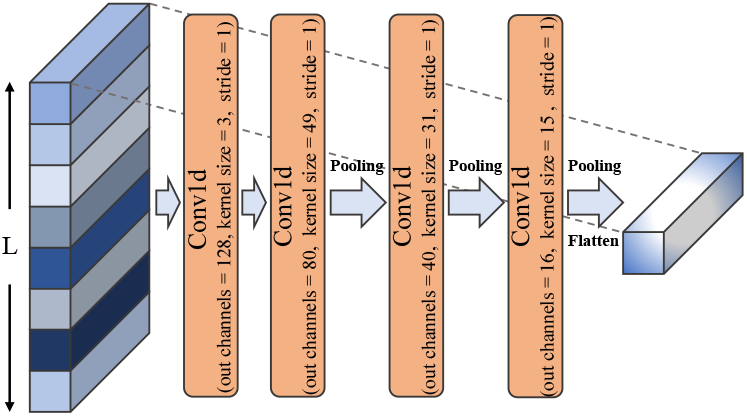
Residue-level feature encoder.

### E. Predictor and Optimization

Given a protein complex pair (ℳ*T, WT*), the graph-level structural feature and multi-perspective sequential features are concatenated to obtain the final representation **P** = [***p***_*graph*_; ***p***_*global*_; ***p***_*context*_; ***p***_*residue*_] ∈ ℝ^6*×d*^. Then, a MLP predictor is used to compute the final representations:

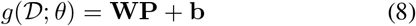

where *θ* is the set of model parameters, *D* denotes the dataset, and **W** and **b** are the predictor weights and bias, respectively.

The objective function is defined as follows:

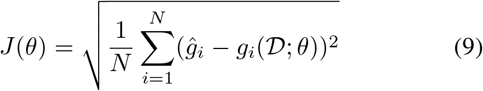

where *N* is the number of instances within the dataset, *θ* represents all the parameters of the model, and *ĝ*_*i*_ and *g*_*i*_ represent the experimental and predicted binding affinity change values for the *i*-th instance, respectively.

**TABLE I.** Performance metrics for the different models on S1131.

| Model | PCC ↑ | RMSE ↓ | MAE ↓ |
| --- | --- | --- | --- |
| FoldX [12] | 0.457 | 2.307 | 1.689 |
| Profile-score [16] | 0.675 | 1.823 | 1.338 |
| BindProfX [17] | 0.738 | 1.701 | 1.247 |
| TopNetTree [4] | 0.850 | 1.550 | 1.138 |
| PerSpect-EL [5] | 0.853 | 1.303 | 0.947 |
| GeoPPI [6] | 0.855 | 1.298 | 0.941 |
| Hom-ML-V2 [18] | 0.857 | 1.279 | 0.939 |
| PTA-SEL [19] | 0.866 | 1.216 | 0.893 |
| PIPR+S2F [13] | 0.866 | 1.210 | 0.886 |
| TransABseq [20] | 0.843 | 1.157 | 0.844 |
| SFM-Net [7] | 0.881 | 1.142 | 0.818 |
| **MPFIL** | **0.917** | **0.983** | **0.718** |

**TABLE II.** Performance metrics for the different models on S2398.

| Model | PCC ↑ | RMSE ↓ | MAE ↓ |
| --- | --- | --- | --- |
| PPA-Pred [21] | 0.65 | 1.672 | 1.138 |
| ProAffiMuSeq [22] | 0.75 | - | - |
| SAAMBE-SEQ [23] | 0.83 | 1.201 | 0.817 |
| GeoPPI [6] | 0.84 | 1.229 | 0.836 |
| PIPR+S2F [13] | 0.86 | 1.190 | 0.810 |
| **MPFIL** | **0.882** | **0.891** | **0.613** |

## IV. Experimental Results and Discussion

### A. Datasets and Evaluation Metrics

Three benchmark datasets are adopted in our experiments:

- *SKEMPI* [8], a dataset of structural and energetic changes associated with mutant protein interactions in heterodimeric protein complexes. A subset of 1,131 non-redundant interface mutations, referred to as S1131 [17].
- *SKEMPI-2*.*0* [9], an expanded version of SKEMPI that reports binding free energy changes upon mutation for structurally resolved PPIs. The subset S2398 [23] is used for prediction, which includes single-chain, single-point, and multi-point mutations, comprising 2,398 instances.
- *MPAD* [10], a membrane protein complex binding affinity database containing experimental affinities of membrane protein–protein complexes and their mutants, together with sequence information, functional annotations, membrane-specific features, and experimental conditions.

Following previous studies, Pearson’s correlation coefficient (PCC) value, root mean squared error (RMSE) value, and mean absolute error (MAE) value are used as evaluation metrics in our experiments. To fairly evaluate the performance of the proposed method, we follow the same strategy as in previous studies [14], [18], splitting the training and testing sets for ten-fold cross-validation experiments.

### B. Performance Comparison

To evaluate the effectiveness of our method, we compare the proposed MPFIL model with several competitive models and state-of-the-art models, including FoldX [12], Profile-score [16], BindProfX [17], TopNetTree [4], PerSpect-EL [5], GeoPPI [6], HCML [18], PTA-SEL [19], PIPR+S2F [13], TransABseq [20], SFM-Net [7], PPA-Pred [21], ProAffiMuSeq [22], and SAAMBE [23]. Tables I and II report various PCC, RMSE, and MAE values of different baseline models and the proposed MPFIL method on the S1131 and S2398 datasets, respectively.

The proposed MPFIL framework has the best performance for ΔΔ*G* prediction. MPFIL achieves the best PCC values of 0.917 and 0.882 on S1131 and S2398, respectively, and also attains the lowest RMSE values of 0.983 and 0.891 on the two datasets. In addition, it obtains the lowest MAE values of 0.718 and 0.613 on S1131 and S2398, respectively. MPFIL demonstrates significant improvements in prediction accuracy by comprehensively incorporating the results achieved on the benchmark datasets. In particular, it outperforms other results in PCC, RMSE, and MAE values by at least 4.0%, 13.9%, and 12.2% on S1131, respectively. It also outperforms other results in such measures by at least 2.5%, 25.1%, and 24.3% on S2398, respectively.

MPFIL shows high effectiveness in exploiting and utilizing the structural-sequential features of protein complexes. Compared with methods that incorporate structural and biological features, including GeoPPI, HCML, and SFM-Net, our method also achieves better results on both datasets. For example, MPFIL outperforms SFM-Net by 4.0% and 13.9% in PCC and RMSE values, respectively, on the S1131 dataset. It out-performs GeoPPI by 5.0% and 27.5% in the same metrics on the S2398 dataset, respectively. Furthermore, compared with sequence-based methods, including PIPR+S2F and SAAMBE-SEQ, the proposed method also achieves better performance values. For instance, our method outperforms SAAMBE-SEQ by 6.2% and 25.8% in terms of PCC and RMSE values, respectively, on the S2398 dataset. The consistent performance improvements of MPFIL on both datasets demonstrate its greater effectiveness in protein complex modeling, multi-level feature capture and exploitation, amino acid interaction learning, and the analysis of mutation-induced structural differences.

### C. Membrane Protein Dataset Evaluation

To further evaluate the effectiveness of the proposed method, in this section, we compare our model with several baseline models on MPAD, a dataset for membrane protein complexes [10]. It is estimated that 20–30% of all proteins in living organisms are membrane proteins, highlighting their critical roles in cellular processes. In comparison with SKEMPI, which primarily consists of soluble proteins, the structural data available for membrane proteins is limited, and structure-based methods tend to exhibit suboptimal performance on such datasets. Therefore, the membrane protein dataset presents a substantial challenge for ΔΔ*G* prediction. Its subset is used in our experiments, comprising 1,040 mutations from 89 proteins [14]. We compare MPFIL with several baseline methods, including several competitive ML methods, structure-based methods (MutaBind2 [24] and SAAMBE-3D [25]), and sequence-based methods (SAAMBE-SEQ [23] and SAAMBE-MEM [14]). Table III reports the PCC, RMSE, and MAE values on the MPAD dataset.

From Table III, our observations can be summarized as follows. The proposed MPFIL method outperforms all other methods for ΔΔ*G* prediction. It shows considerable advancement in PCC, achieving the highest value of 0.713 and outperforming baseline models by at least 11.4%. It also achieves the lowest RMSE value of 0.729, surpassing sequence-based methods and ML methods by at least 7.7% and 12.1%, respectively. Furthermore, MPFIL attains the lowest MAE value of 0.545, surpassing other baseline models by at least 5.8%. The proposed method maintains its superior performance even on the membrane protein dataset with limited structural information.

**TABLE III.**
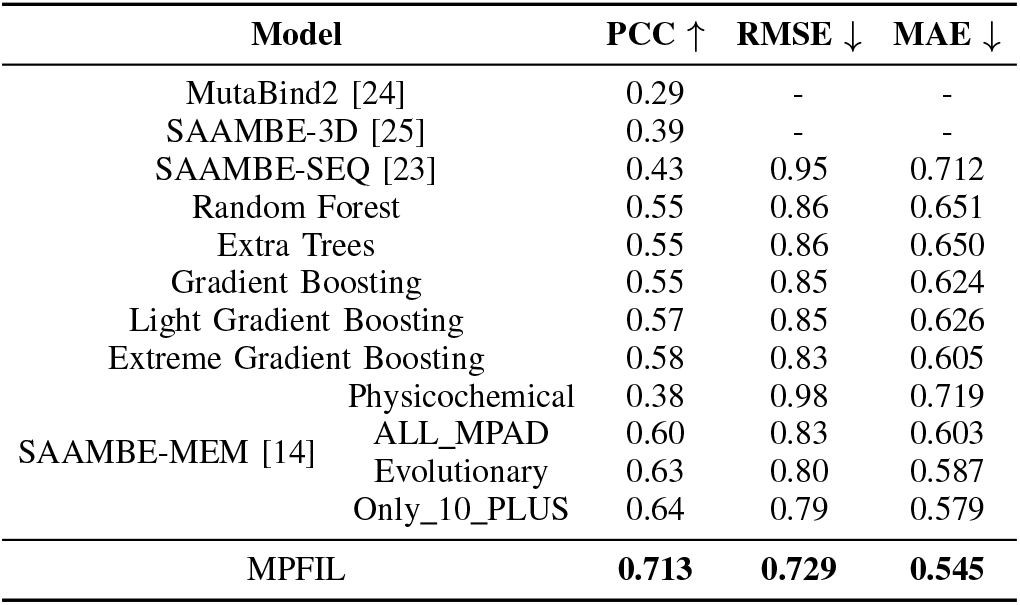
Performance metrics for different models on MPAD.

The comparative results on the MPAD dataset demonstrate the effectiveness and universality of MPFIL in the domain of membrane protein ΔΔ*G* prediction.

### D. Ablation Study

To analyze the contributions and effects of different components of our framework, we conducted ablation experiments. We experiment with MPFIL and its four variants: the model without graph-level structural feature (i.e., MPFIL w/o *structure*), the model without global level differential feature (i.e., MPFIL w/o *global*), the model without full-sequence context-level feature (i.e., MPFIL w/o *context*), and the model without residue-level sequential feature (i.e., MPFIL w/o *residue*). The results are presented in Table IV, based on which, our observations can be summarized as follows:

When the graph-level structural feature is removed, the prediction performance of the corresponding variant is substantially degraded, particularly on the MPAD dataset. Specifically, MPFIL outperforms the variant (w/o *structure*) by 16.5%, 11.8%, and 12.6% in PCC, RMSE, and MAE, respectively, on the MPAD dataset. It also attains performance improvements of at least 5.2%, 7.3%, and 9.5% in the mean values of these metrics. The results suggest that the graph-level structural feature encoding module possesses the capacity to accurately describe and capture the impact of mutations on protein structures, thereby enabling the effective learning of the structural changes and critical differential information in complexes caused by mutations.

In comparison with the variants that do not utilize multi-perspective sequential features, MPFIL achieves varying degrees of improvement in prediction performance. In particular, it outperforms the variant (w/o *residue*) by at least 2.9%, 7.5%, and 9.6% in the mean metrics of PCC, RMSE, and MAE values, respectively. A comparative analysis of MPFIL with the variants (w/o *global*) and (w/o *context*) reveals that the proposed method achieves comparable improvements across all metrics on three benchmark datasets. The experimental results suggest that each level feature within the multi-perspective sequential features plays a crucial role in accurately predicting ΔΔ*G*. We speculate the reason for the performance gain is that the global level differential feature encompasses the differential information of the protein sequences at the global level; the residue-level sequential feature accurately captures the medium-range and short-range dependencies within the protein sequence, particularly in regions proximal to the mutation site; and the full-sequence context-level feature captures and learns long-range dependencies, semantic relationships, and interactions between amino acids across the entire protein complex sequence.

### E. Distribution Comparisons

The performance of the proposed framework is further evaluated through comparative analyzes of the distributions of experimental and predicted ΔΔ*G* values from various perspectives. Figure 4 presents a density scatter plot that compares the predicted values with experimentally determined values on the benchmark datasets. Figure 5 presents the relationship between the experimental values and the predicted values. The data points are categorized into three distinct regions based on the experimental ΔΔ*G*, thereby illustrating the model’s behavior across the specified ranges. Figure 6 presents detailed distribution comparisons across three benchmark datasets.

**Fig 4.**
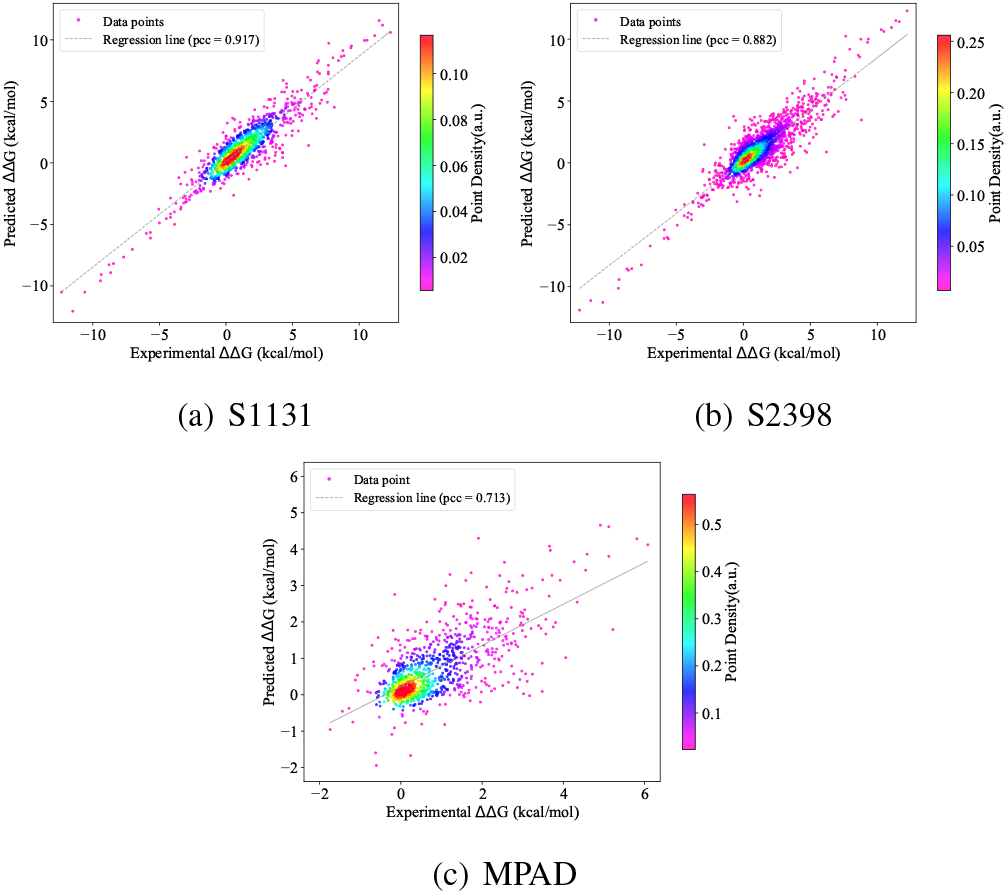
Prediction performance of MPFIL on the benchmark datasets.

**Fig 5.**
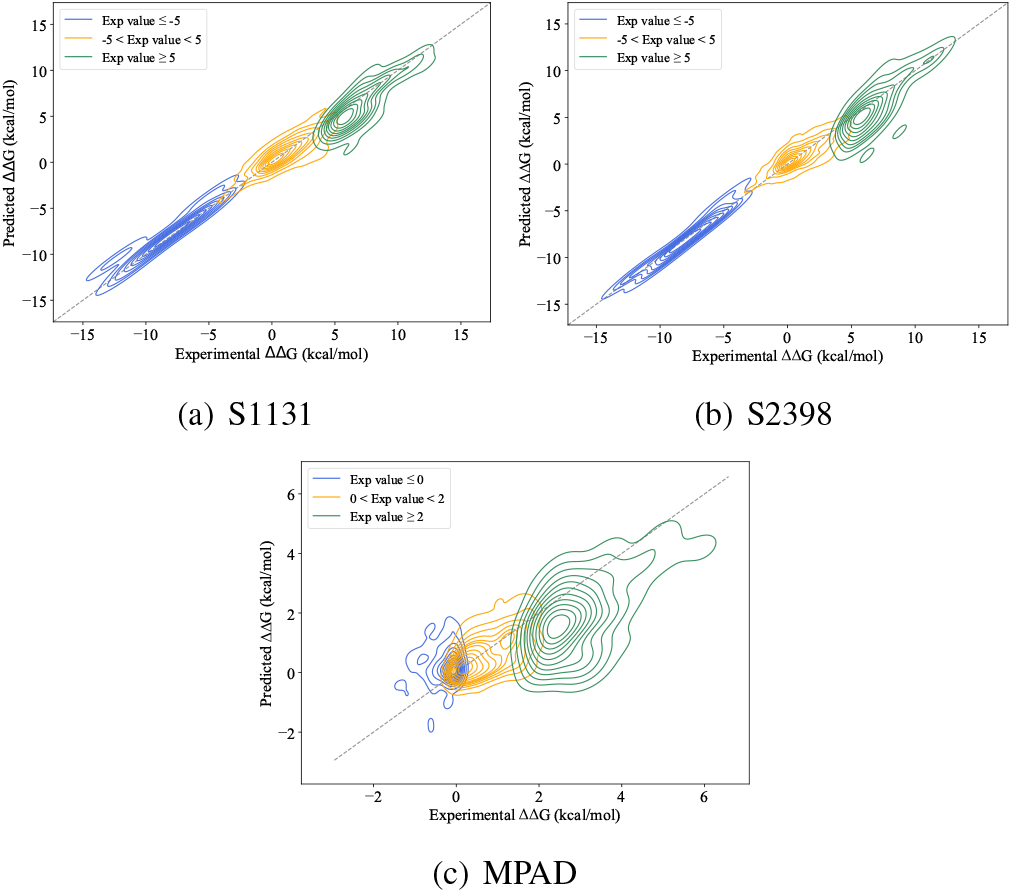
The predicted ΔΔ*G* values against the experimental ΔΔ*G* values across different distinct ranges on the benchmark datasets.

**Fig 6.**
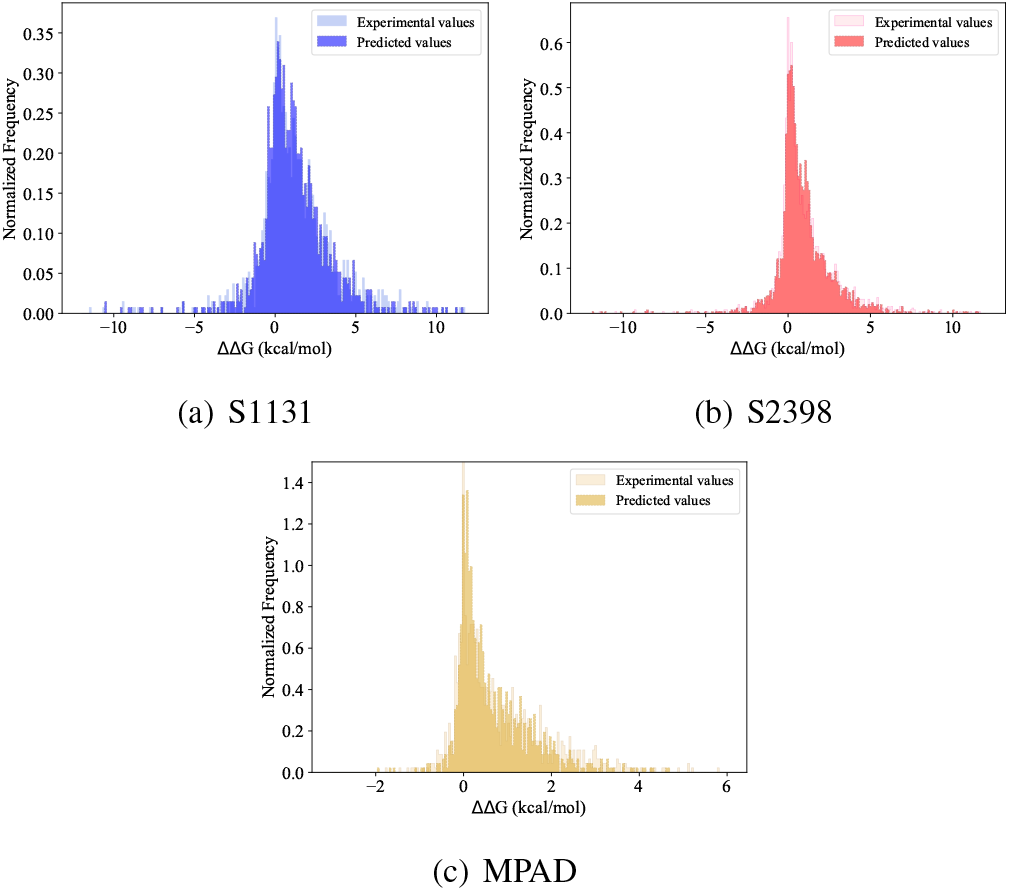
Distribution comparisons of ΔΔ*G* values on the benchmark datasets.

In Figure 4, MPFIL demonstrates its excellent prediction performance on all datasets. As shown in Figure 4(a) and (b), the majority of data points are concentrated around the diagonal, indicating the satisfactory prediction performance of the proposed method for ΔΔ*G* values within the central range. In Figure 4(c), the predicted sample points are more concentrated along the regression line in the central region. As shown in Figure 6, the prediction results closely match the distributions of experimental values across three benchmark datasets. The comparative results demonstrate that our method can accurately predict ΔΔ*G* values and exhibit excellent capability for protein modeling and feature extraction.

## V. Conclusion and Future Work

In this paper, we propose a novel DL-based multi-level protein feature integrated learning framework and demonstrate its effectiveness in predicting mutation-induced binding affinity changes. Our framework effectively integrates information-rich structural features with multi-perspective sequential features of protein complexes to enable an accurate description of mutation behavior and ΔΔ*G* prediction. Experiments on three benchmark datasets demonstrated that: our method effectively extracts protein structural features and learns structural changes and differential information induced by mutations; multi-level sequential feature encoding enables effective learning of long- and mid-range dependencies across the entire protein sequence, intrinsic interactions between residues, global semantic relationships, and comprehensive biological regulatory information. On all datasets, the proposed method consistently outperforms competitive baseline methods, demonstrating its broad applicability to ΔΔ*G* prediction for both soluble and membrane proteins.

**TABLE IV.** Mtrics values of model with different components on the benchmark datasets.

| Model | PCC ↑ | | | | RMSE ↓ | | | | MAE ↓ | | | |
| --- | --- | --- | --- | --- | --- | --- | --- | --- | --- | --- | --- | --- |
|  | S1131 | S2398 | MPAD | Mean | S1131 | S2398 | MPAD | Mean | S1131 | S2398 | MPAD | Mean |
| MPFIL w/o *structure* | 0.903 | 0.870 | 0.612 | 0.795 | 1.061 | 0.922 | 0.827 | 0.937 | 0.828 | 0.622 | 0.624 | 0.691 |
| MPFIL w/o *global* | 0.904 | 0.877 | 0.696 | 0.826 | 1.056 | 0.904 | 0.748 | 0.903 | 0.803 | 0.614 | 0.561 | 0.659 |
| MPFIL w/o *context* | 0.907 | 0.872 | 0.682 | 0.820 | 1.039 | 0.915 | 0.762 | 0.905 | 0.796 | 0.619 | 0.572 | 0.662 |
| MPFIL w/o *residue* | 0.891 | 0.867 | 0.681 | 0.813 | 1.127 | 0.931 | 0.760 | 0.939 | 0.869 | 0.636 | 0.571 | 0.692 |
| **MPFIL** | **0.917** | **0.882** | **0.713** | **0.837** | **0.983** | **0.891** | **0.729** | **0.868** | **0.718** | **0.613** | **0.545** | **0.625** |

Beyond the work presented here, systematic analysis of disease-associated mutations is clearly of interest for future research. We plan to explore the application of the proposed method to variants linked to cancer and neurodegenerative diseases, thereby facilitating the development of precision medicine and preventive healthcare strategies.

## REFERENCES

[1] L. Hu, X. Wang, Y.-A. Huang, P. Hu, and Z.-H. You, “A survey on computational models for predicting protein–protein interactions,” Briefings in bioinformatics, vol. 22, no. 5, p. bbab036, 2021.

[2] F. Cheng, J. Zhao, Y. Wang, W. Lu, Z. Liu, Y. Zhou, W. R. Martin, R. Wang, J. Huang, T. Hao et al., “Comprehensive characterization of protein–protein interactions perturbed by disease mutations,” Nature genetics, vol. 53, no. 3, pp. 342–353, 2021.

[3] C. H. Rodrigues, D. E. Pires, and D. B. Ascher, “mmcsm-ppi: predicting the effects of multiple point mutations on protein–protein interactions,” Nucleic Acids Research, vol. 49, no. W1, pp. W417–W424, 2021.

[4] M. Wang, Z. Cang, and G.-W. Wei, “A topology-based network tree for the prediction of protein–protein binding affinity changes following mutation,” Nature Machine Intelligence, vol. 2, no. 2, pp. 116–123, 2020.

[5] J. Wee and K. Xia, “Persistent spectral based ensemble learning (perspect-el) for protein–protein binding affinity prediction,” Briefings in Bioinformatics, vol. 23, no. 2, p. bbac024, 2022.

[6] X. Liu, Y. Luo, P. Li, S. Song, and J. Peng, “Deep geometric representations for modeling effects of mutations on protein-protein binding affinity,” PLoS computational biology, vol. 17, no. 8, p. e1009284, 2021.

[7] C. Liu, S. Cai, T. Pan, H. Ogata, J. Song, and T. Akutsu, “Sfm-net: Selective fusion of multiway protein feature network for predicting binding affinity changes upon mutations,” Journal of Chemical Information and Modeling, 2025.

[8] I. H. Moal and J. Fernández-Recio, “Skempi: a structural kinetic and energetic database of mutant protein interactions and its use in empirical models,” Bioinformatics, vol. 28, no. 20, pp. 2600–2607, 2012.

[9] J. Jankauskaitć, B. Jiménez-García, J. Dapkănas, J. Fernández-Recio, and I. H. Moal, “Skempi 2.0: an updated benchmark of changes in protein–protein binding energy, kinetics and thermodynamics upon mutation,” Bioinformatics, vol. 35, no. 3, pp. 462–469, 2019.

[10] F. Ridha, A. Kulandaisamy, and M. M. Gromiha, “Mpad: A database for binding affinity of membrane protein–protein complexes and their mutants,” Journal of Molecular Biology, vol. 435, no. 14, p. 167870, 2023.

[11] M. Li, F. L. Simonetti, A. Goncearenco, and A. R. Panchenko, “Mutabind estimates and interprets the effects of sequence variants on protein– protein interactions,” Nucleic acids research, vol. 44, no. W1, pp. W494– W501, 2016.

[12] J. Delgado, L. G. Radusky, D. Cianferoni, and L. Serrano, “Foldx 5.0: working with rna, small molecules and a new graphical interface,” Bioinformatics, vol. 35, no. 20, pp. 4168–4169, 2019.

[13] Y. Xue, Z. Liu, X. Fang, and F. Wang, “Multimodal pre-training model for sequence-based prediction of protein-protein interaction,” in Machine Learning in Computational Biology. PMLR, 2022, pp. 34–46.

[14] P. Rimal, S. K. Panday, W. Xu, Y. Peng, and E. Alexov, “Saambe-mem: a sequence-based method for predicting binding free energy change upon mutation in membrane protein–protein complexes,” Bioinformatics, vol. 40, no. 9, p. btae544, 2024.

[15] A. Elnaggar, M. Heinzinger, C. Dallago, G. Rehawi, Y. Wang, L. Jones, T. Gibbs, T. Feher, C. Angerer, M. Steinegger et al., “Prottrans: Toward understanding the language of life through self-supervised learning,” IEEE transactions on pattern analysis and machine intelligence, vol. 44, no. 10, pp. 7112–7127, 2021.

[16] J. R. Brender and Y. Zhang, “Predicting the effect of mutations on protein-protein binding interactions through structure-based interface profiles,” PLOS Computational Biology, vol. 11, no. 10, pp. 1–25, 10 2015.

[17] P. Xiong, C. Zhang, W. Zheng, and Y. Zhang, “Bindprofx: assessing mutation-induced binding affinity change by protein interface profiles with pseudo-counts,” Journal of molecular biology, vol. 429, no. 3, pp. 426–434, 2017.

[18] X. Liu, H. Feng, J. Wu, and K. Xia, “Hom-complex-based machine learning (hcml) for the prediction of protein–protein binding affinity changes upon mutation,” Journal of chemical information and modeling, vol. 62, no. 17, pp. 3961–3969, 2022.

[19] L. Xiang and K. Xia, “Persistent tor-algebra based stacking ensemble learning (pta-sel) for protein-protein binding affinity prediction,” in ICLR 2022 Workshop on Geometrical and Topological Representation Learning, 2022.

[20] C.-F. Li, Z. Yan, F. Ge, X. Yu, J. Zhang, M. Zhang, and D.-J. Yu, “Transabseq: A two-stage approach for predicting antigen–antibody binding affinity changes upon mutation based on protein sequences,” Journal of Chemical Information and Modeling, vol. 65, no. 10, pp. 5188–5204, 2025.

[21] K. Yugandhar and M. M. Gromiha, “Protein–protein binding affinity prediction from amino acid sequence,” Bioinformatics, vol. 30, no. 24, pp. 3583–3589, 2014.

[22] S. Jemimah, M. Sekijima, and M. M. Gromiha, “Proaffimuseq: sequence-based method to predict the binding free energy change of protein–protein complexes upon mutation using functional classification,” Bioinformatics, vol. 36, no. 6, pp. 1725–1730, 2020.

[23] G. Li, S. Pahari, A. K. Murthy, S. Liang, R. Fragoza, H. Yu, and E. Alexov, “Saambe-seq: a sequence-based method for predicting mutation effect on protein–protein binding affinity,” Bioinformatics, vol. 37, no. 7, pp. 992–999, 2021.

[24] N. Zhang, Y. Chen, H. Lu, F. Zhao, R. V. Alvarez, A. Goncearenco, A. R. Panchenko, and M. Li, “Mutabind2: predicting the impacts of single and multiple mutations on protein-protein interactions,” Iscience, vol. 23, no. 3, 2020.

[25] S. Pahari, G. Li, A. K. Murthy, S. Liang, R. Fragoza, H. Yu, and E. Alexov, “Saambe-3d: predicting effect of mutations on protein– protein interactions,” International journal of molecular sciences, vol. 21, no. 7, p. 2563, 2020.

